# Computer vision guided rapid and precise automated cranial microsurgeries in rodents

**DOI:** 10.1101/2024.09.03.611036

**Authors:** Zahra S Navabi, Ryan Peters, Beatrice Gulner, Arun Cherkkil, Eunsong Ko, Farnoosh Dadashi, Jacob O Brien, Michael Feldkamp, Suhasa B Kodandaramaiah

**Affiliations:** Department of Mechanical Engineering, University of Minnesota, Twin Cities, MN; Department of Computer Science and Engineering, University of Minnesota, Twin Cities, MN; Department of Biomedical Engineering, University of Minnesota, Twin Cities, MN; Department of Neuroscience, University of Minnesota, Twin Cities, MN

## Abstract

Neuroscientists employ various experimental procedures to interface with the brain to study and perturb the neural activity during behavior. A common procedure that allows such physical interfacing is cranial microsurgery, wherein small to large craniotomies are performed in the overlying skull for insertion of neural interfaces or implantation of optically clear windows for long-term cranial observation. Performing craniotomies is, however, a skilled task that requires significant time and practice and further needs to be carried out precisely to ensure that the procedure does not cause damage to the underlying brain and dura. Here, we present a computer vision-guided craniotomy robot (CV-Craniobot) that utilizes machine learning to accurately estimate the dorsal skull anatomy from optical coherence tomography (OCT) images. Instantaneous information of the skull morphology is used by a robotic mill to rapidly and precisely remove the skull from a desired craniotomy location. We show that the CV-Craniobot can perform small (2 - 4 mm diameter) craniotomies with near 100% success rates within 2 minutes and large craniotomies encompassing most of the dorsal cortex in less than 5 minutes. Thus, the CV-Craniobot enables rapid and precise craniotomies, significantly reducing surgery time as compared to human practitioners and eliminating the need for long training.

## INTRODUCTION

Much of neuroscience that uses rodent models such as mice and rats require access to the brain by removing the overlying skull to insert neural probes for recording, optical imaging, and perturbation of neural activity. In recent years, the need to perform simultaneous multi-area recordings has led neuroscientists to increasingly remove large sections of the dorsal skull to allow optical imaging and access to penetrating neural probes for targeting deep brain regions. The advent of ultra-widefield imaging systems such as the two-photon mesoscopes [1] has prompted research groups to develop surgical approaches to perform very large craniotomies across the dorsal cortex [2], [3], [4], [5] and the cerebellar cortex [6]. Once removed, the skull is typically replaced with either curved glass windows [2] or 3D-printed polymer skulls that conform to the morphology of the removed skull [7], [8], [9], [10], [11], [12]. Such approaches have also been adapted for implanting large cranial windows with access ports to insert multiple neural recording devices [5].

While these approaches provide unprecedented capability and flexibility to access and dissect neural function, the challenging craniotomy procedures needed to implant such large cranial windows have precluded widespread adoption. Even more widely adopted cranial window implantation techniques for imaging smaller fields of view [13], [14] require practice and have variable learning curves depending on individual surgeons. Thus, standardized automation of skull removal would be a highly useful tool for neuroscience research.

An accurate estimate of the skull thickness is necessary for automated bone removal. Various approaches have addressed this issue; some use destructive methods to find the optimal milling depth automatically. These include systems that use change in the electrical impedance sensed by the tip of the driller to specify the type of tissue in contact with the drill tip [15], systems that use mechanical impedance of the tissue under the driller tip [16], or drills that use vibration and sound [17] to detect reaching the brain during milling. However, these approaches are prone to false positives when hitting blood vessels in the bone [15]. Furthermore, they are difficult to implement on intricate rodent skulls [16]. Thus, previous approaches have relied on substantial user inputs during robotic milling and require repeated runs to shave off enough material from the skull surface to break off the bone [18]. To date, there have not been any approaches that do not require human intervention during automated craniotomies.

Here, we describe the CV-Craniobot, a computer vision-guided craniotomy robot that combines non-destructive imaging of live mouse skulls with robotic milling to fully automate craniotomies in mice. The CV-Craniobot utilizes an optical coherence tomography (OCT) scanner to image through the skull. Machine learning models are trained to accurately reconstruct the skull’s dorsal and ventral surface from the OCT images. The machine learning model’s reconstructions of the dorsal and ventral surface of the skull can be used to perform rapid and precise robotic removal of bone, with a near 100% success rate at a fraction of the time taken to perform manual craniotomies.

## RESULTS

### CV-Craniobot operation and architecture

The operation of the CV-Craniobot is shown in **Figure 1A**. First, the mouse skull is affixed in a custom-made stereotax integrated within the robot; then, it is scanned using an optical coherence tomography (OCT) scanner. An OCT system uses low-coherence interferometry to image through the different layers of scattering material [19] [20]. Here, we use an OCT system with a center wavelength of 1310 nm and a bandwidth of 60 nm. This enables us to image non-destructively through the bone tissue and obtain a near-instantaneous 3D reconstruction of both the dorsal and ventral skull surfaces. To estimate the pixels containing the bone tissue, CV- Craniobot employs a machine learning model (U-Net), which is trained to segment the bone tissue in the B-scans (X-Z cross-sections of the 3D OCT scan). The segmented images are then used to identify the top edge and bottom edges of the skull tissue. Using the extracted edges on all B-scans, the skull’s top and bottom 3D boundaries- the dorsal and ventral surfaces of the mouse skull- are reconstructed. Once the surfaces are estimated, this information and the user input milling path guide a robotic drill to automatically perform bone milling using the estimated dorsal or ventral skull surface as a reference.

**Figure 1:**
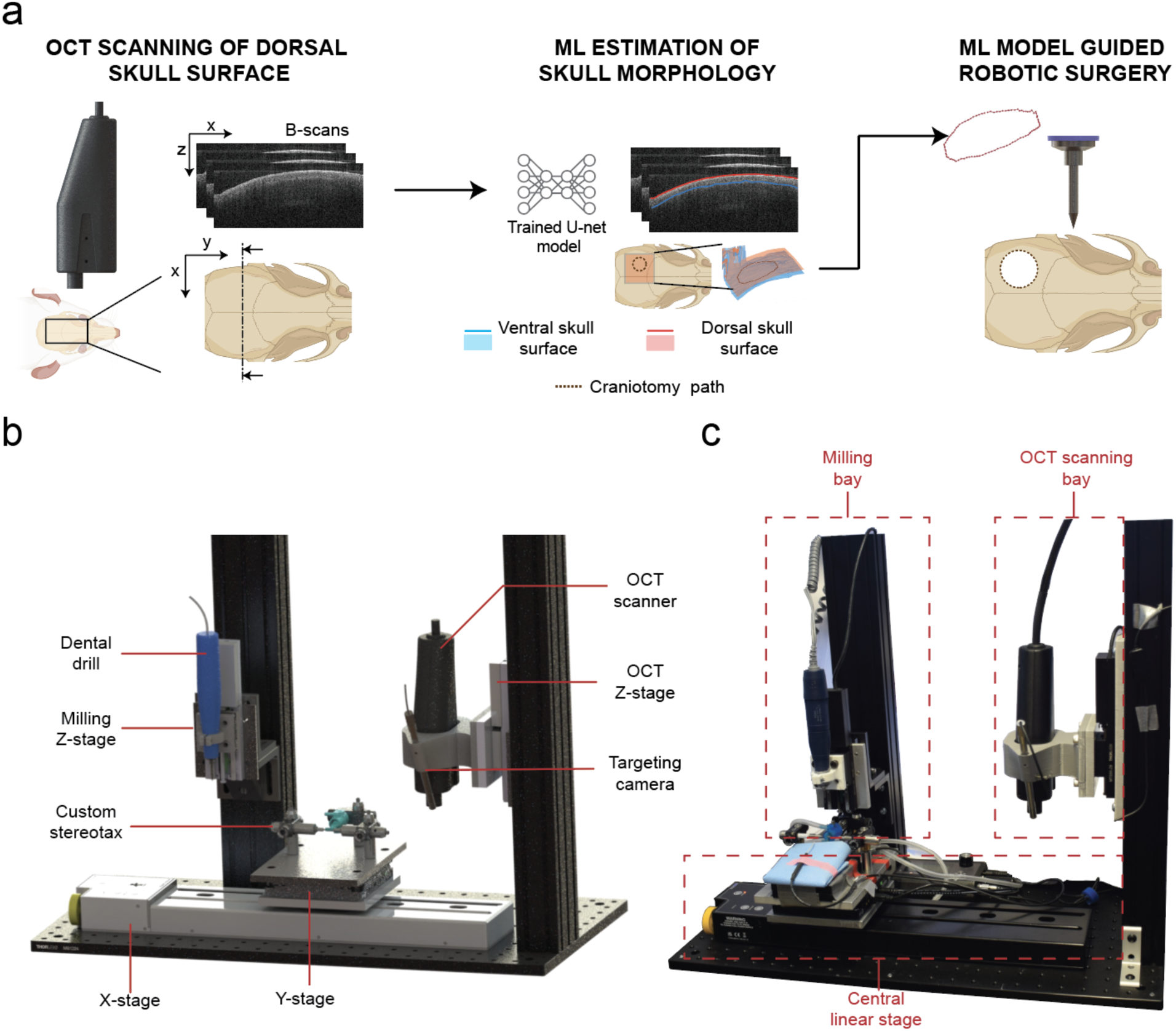
**CV-Craniobot working principle and hardware architecture**: **(a)** Workflow: The mouse skull is imaged using an OCT scanner, an ML model estimates the dorsal and ventral skull surfaces from the B-scans of the OCT image; a robotic mill executes automated bone removal along the computed craniotomy path. **(b)** Computer-aided design (CAD) rendering of the CV-Craniobot hardware. **(c)** Photograph of the CV-Craniobot with core elements highlighted.

To execute this operation principle, we established the CV-Craniobot shown in **Fig. 1B**. The CV- Craniobot consists of three major components: First, a central linear stage on which a custom stereotax is installed. Second, the OCT scanning bay, where the OCT scanner is mounted on an adjustable z-stage, allows the OCT scanner to focus properly on the mouse skull. Third, the milling bay, which is equipped with an electrical dental drill mounted on a Z-axis linear stage, adjusts the height of the driller with respect to the skull, while the central linear stage controls the lateral position of the driller with respect to the skull. The linear stage transports the mouse affixed to the stereotax between an OCT scanning bay and the craniotomy milling bay.

### OCT imaging and ML estimation of the skull surface

OCT scans are known to be prone to different types of distortions [21], [22], [23], which need to be corrected before they can be used for accurately estimating skull morphology. These distortions include non-linear scanning distortions that can occur in the axial (Z axes) direction, non-telecentric distortions that cause images to be warped laterally (XY plane), and optical distortions that cause distortion in the axial direction when imaging through materials with different refractive index. Imaging a flat glass coverslip using the OCT scanner revealed negligible non-linear axial scanning distortion (**Supplementary Fig. 1**). We imaged a modeling clay sample imprinted with a mesh pattern with an approximate spacing of 125 µm to determine lateral image warping. Reconstruction of the raw OCT images revealed both tangential and radial distortions in the lateral direction (X-Y plane, **Fig. 2B, *left***). Image distortion in both X and Y axes increased in the lateral areas of the OCT’s field of view (FOV), resulting in an average of 13 pixels distortion in the most lateral X-lines and an average of 8 pixels distortion in the Y-lines. The distortion was measured as the root mean square error between the detected lines in each cross-section and the reference (undistorted) line corresponding to the detected line (**see Methods**). The division model for radial distortion and the Brown-Conrady model for tangential distortion were employed to represent and correct for distortions in the X-Y plane (See **Methods**). After correction, images had a measured distortion of around 1 pixel in both directions (**Fig. 2C** and **D**). Images of the mesh after distortion correction are shown in **Figure 2B, *right panel***.

**Figure 2:**
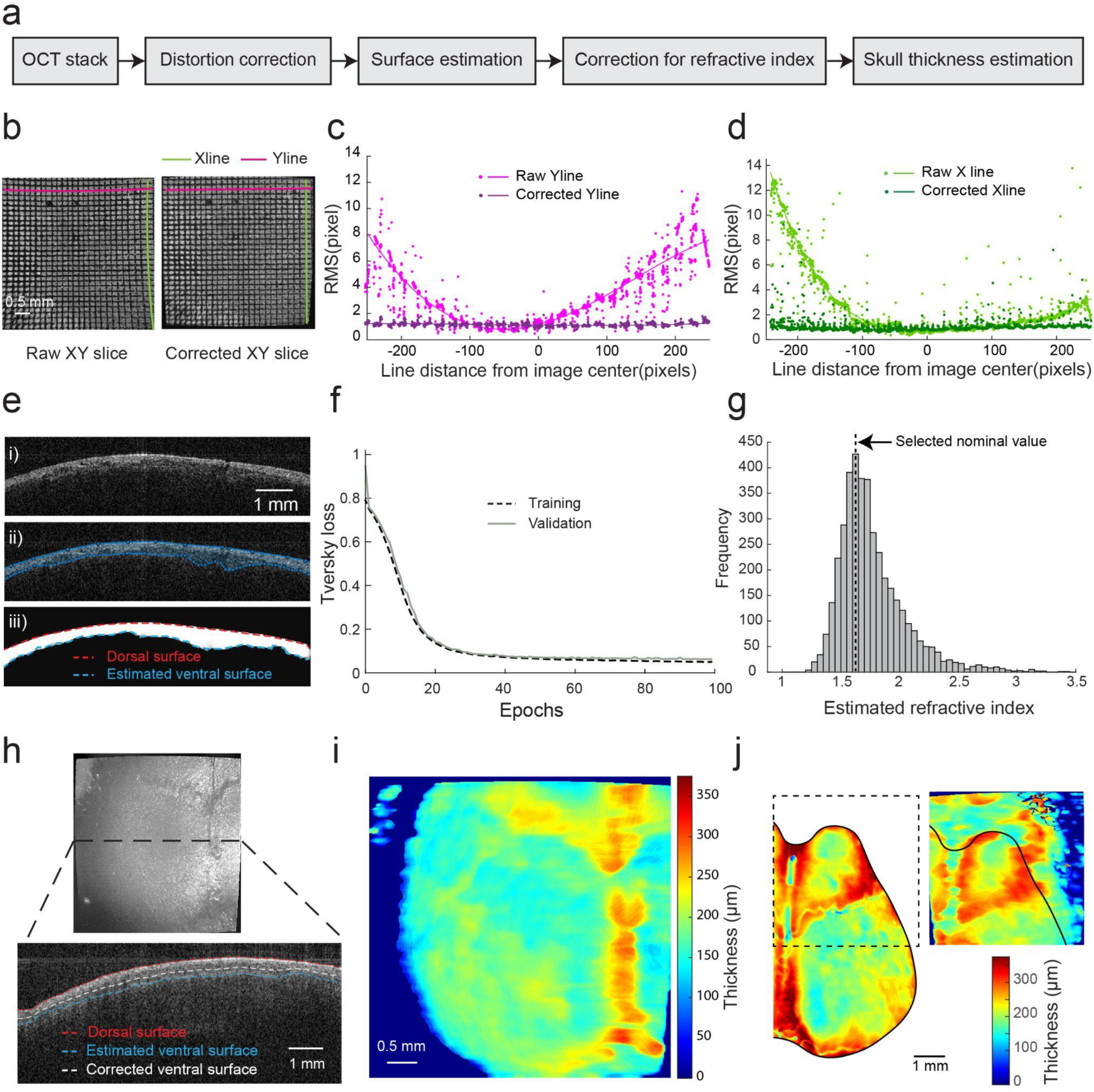
Rapid, accurate estimation of skull morphology from OCT image scan: (a) Steps involved in estimating skull morphology from OCT image scan **(b)** Reconstructed top view image of a modeling clay sample imprinted with mesh pattern before and after non-telecentric distortion correction. **(c)** Root Mean Square Error (RMSE) between each Y-line and the reference line at that position for both raw and corrected scans. **(d)** RMSE between the raw and corrected X-lines. **(e)** U-Net Model training i. Representative average B-scan image from OCT scan of a mouse skull. ii. B-Scan image with manually annotated dorsal and ventral skull surfaces. iii. Trained U-model estimate of the dorsal and ventral surfaces. **(f)** Tversky loss plot for training and validation. **(g)** Measured refractive index (n= 8 mice) (See **Supplementary Note 1**). **(h)** *Top:* Reconstructed top view of the mouse’s dorsal skull from OCT scan, *bottom:* B-scan cross-section with estimated dorsal and ventral surfaces by U-Net model and the refractive index-corrected ventral surface. **(i)** A pseudo-color map of the skull thickness of the specimen is shown in **h**. **(j)** Comparison between the microcomputer tomography (μCT) scan measurement (left) and the estimated thickness using OCT scans in the same area.

Once the images were corrected for non-telecentric distortions, we proceeded to identify the dorsal and ventral skull surfaces from the cross-sectional images acquired from the OCT. The low coherence IR light passing through the dorsal skull surface from air creates a clear contrast in the cross-section image, whereas the ventral surface boundary with the brain tissue underneath was less discernible (**Fig. 2D.i**). To accurately reconstruct the two surfaces, we trained a U-Net model [24], a convolutional neural network model widely used for biomedical image segmentation. The model was trained using 300 B-scans of live mouse skulls, in which the skull bone area was manually annotated (**Fig. 2D.ii**)(See **Methods** for details). Our results demonstrated good generalization to out-of-fold samples, with training and validation accuracies of 0.995 ± 0.001 and 0.994 ± 0.003 (**Fig. 2E**) and Intersection over Unions (IOUs) of 0.908 ± 0.021 and 0.886 ± 0.043 respectively (**Supplementary Figure 2**). The skull’s dorsal and ventral surfaces were then reconstructed using the top and bottom boundaries of all segmented B-scans using image processing methods (**Fig. 2D.iii**).

An OCT system determines scattering depth by measuring the difference in travel time between light returning from the sample and the reference arm. Due to the slower speed of light in bone compared to air, the light takes longer to return from the ventral surface of the skull, causing the ventral surface to appear deeper in OCT images than it actually is [25], [26]. To correct this, the refractive index of the mouse skulls bone tissue was measured (**Fig. 2F**, See also **Supplementary Fig. 3** and **Supplementary Note 1**). The ventral surface estimated by the U-Net model was corrected for the refractive index distortion using a nominal refractive index value of 1.6 (**Fig. 2G**). **Figure 2I** shows a pseudo-color image of the estimated thickness of the skull from the OCT image shown in **Figure 2H**. Consistent with previous work [27], we found that the average skull thickness in the parietal bone region was mostly even, with smooth surfaces defining both the dorsal and ventral skull surfaces, resulting in thickness ranging from 100 to 650 µm[7]. Further, the thickness of the skull increased in the areas immediately surrounding the bone sutures, with the highest thickness measured at the midline suture. The skull thickness images allowed distinguishing the Bregma and Lambda landmarks as well as the bone sutures. Pseudo-color images of a skull generated first with the OCT scanner, followed by µCT images, are shown in **Figure 2J**, showing agreement in the estimation of skull thickness across these modalities.

### Image-guided milling using dorsal skull surface as reference

To evaluate whether we could use the estimated dorsal and ventral skull surfaces for robotic milling, the CV-Craniobot was used to engrave the University of Minnesota logo on the eggshells with increasing engraving depth (**Fig. 3A, Supplementary Video 1**). Eggshells provided a reference surface mimicking the curved surface of the dorsal skull while providing an optically opaque and rigid surface. The robotic milling provided sufficient machining resolution to visually discriminate differences in milling depth, indicating that the surface reconstructed from the OCT images was accurate and suitable for robotic milling. Post-milling scans of the engraved logos were acquired both with a μCT scanner and the OCT scanner to evaluate the milling depth after the procedure (**Fig. 3B** and **C**). Qualitatively, uniform milling depth was observed along the entire length of the milling path. The average depth of milling along the whole milled length was calculated. A clear linear relationship was observed between the milling depth and the commanded depth (R^2^=0.91) (**Fig. 3D**). In general, the milling depth was slightly lesser than the commanded milling depth, possibly due to a combination of system calibration offset and incomplete removal of material along the periphery of the milling tool.

**Figure 3:**
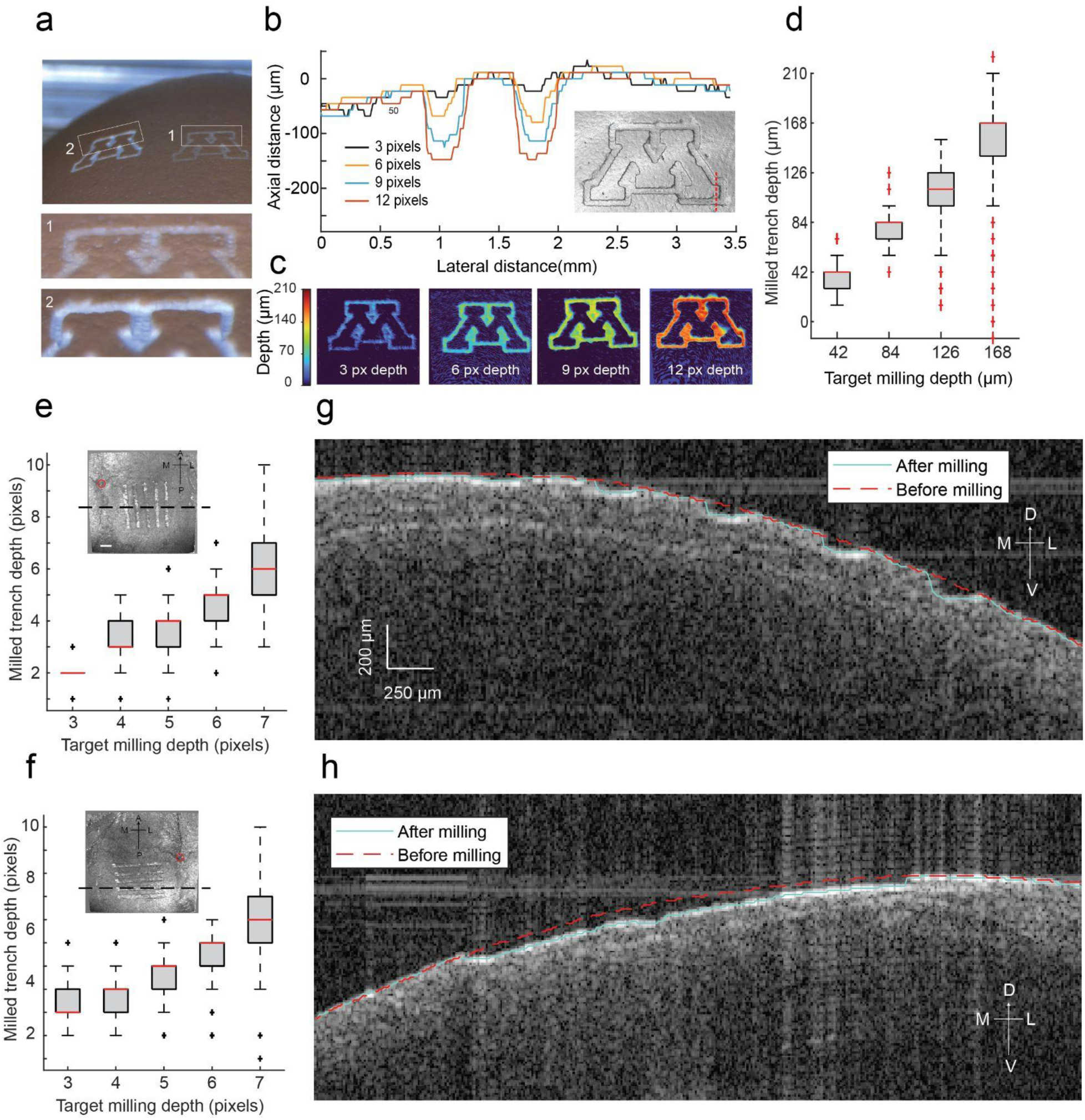
Robotic milling using the estimated dorsal surface as reference: (a) Photograph of the surface of an eggshell engraved with the University of Minnesota logo using the CV-Craniobot. Insets 1 and 2 show close-up images of the engravings milled to depths of milling operations with depths of 42 and 168 µm, respectively. **(b)** Surface profiles of the bottom left corner logo were obtained via µCT scanning of the eggshell surface after engraving the logo on it. Inset: Reconstructed µCT scan top view of the 6 pixels (84 µm) deep logo engraving. The red dashed line indicates the cross-section of the surface shown in the plot. **(c)** Pseudocolor plots of the engraving depth. **(d)** Milled trench depth along the milling path for different target milling depths, as measured using the OCT scanner. **(e)** Milled trench depths along the vertical paths for different target milling depths. *Inset:* Top-view image of the OCT scanner’s FOV after milling. The red circle indicates Bregma. The dashed lines show the cross-section displayed in +**g**. **(f)** Milling depths along the horizontal paths for different target milling depths. *Inset:* Top-view image of the OCT scanner’s FOV after milling. The red circle indicates Bregma. The dashed lines show the cross-section displayed in **h**. **(g)** Coronal section of the mouse skull along the dashed line indicated in **e**. **(h)** Coronal cross-section of the skull along the dashed line indicated in **f**. A - Anterior, P - posterior, M - Medial, L - Lateral, D - Dorsal, V - Ventral.

Next, similar acute experiments were performed on anesthetized mice. CV-Craniobot was used to scan and estimate the dorsal skull surface of a mouse, which was then used as a reference to mill trenches of varying depths in the anteroposterior and mediolateral directions (**Fig. 3E** and **F)**. The CV-Craniobot was programmed to mill 2.2 mm long lines on the parietal bone 0.8 - 3 mm lateral and 0.2 - 2 mm posterior to Bregma (**Fig. E** and **F**) with depths increasing from 3 pixels (42 µm) to 7 pixels (98 µm) along the axial direction of the OCT scan with respect to the dorsal skull surface. Similar to the milling experiments on eggshells, milling the dorsal skull surface along both the anteroposterior and medio-lateral directions resulted in linear tracking of commanded milling depth by milled trench depth. Post-milling scanning of the skull surface revealed clear trenches (**Fig. 3G** and **H**). The comparison between the OCT images taken before and after the milling operations clearly indicated the removal of bone along the trenches (**Fig. 3G** and **H).** Measuring the depth of the milled trenches and comparing with the commanded depth of milling, we again found that there was a clear linear trend (**Fig. 3E** and **F**, R^2^ =0.71 for vertical and R^2^ =0.74 for horizontal lines), with the measured milled depth being ∼14 µm (1 pixel) less than the commanded depth of milling. The high variability in the deeper trenches is partly due to the higher curvature of the skull at the edges and the fixed orientation of the driller. This causes one side of the milled trench to be slightly deeper inside the bone while the other is shallower than the targeted depth.

Having established that the estimated dorsal skull surface could be used for robotic milling of eggshells and live anesthetized mice, we next performed experiments to determine whether the estimated ventral surface could be used as a reference for computerized milling. The CV- Craniobot was programmed to mill down to various depths, leaving a programmed skull thickness above the ventral surface. Circular craniotomies of 2mm diameter were performed on four mice to quantify the accuracy of this milling operation. The robotic mill was commanded to remove bone, leaving a 70 µm (5 pixels in OCT image coordinates) thick layer of bone tissue above the ventral skull surface. The craniotomies were performed, centered at ∼1.8 mm anterior and ∼1.4 mm lateral to Bregma (**Fig. 4A**). Cross-sectional OCT images illustrated that we were indeed able to reach the desired depth of milling, with the median thickness of the remaining bone across four craniotomies being 82 µm with interquartile range (IQR) in 71µm-101µm (6 pixels-IQR: 5-7, **Fig. 4F-left, Supplementary Fig. 4A**). The same procedure (n=4 mice) was also performed with a target remaining thickness of 98µm (7 pixels). This time, the craniotomies were performed at different positions with respect to the Bregma to sample the robot’s performance in different skull regions. After milling along the programmed path, the OCT-measured resultant thickness had a median value of 106 µm with IQR: 92 µm-122µm (7 pixels IQR:6-8, **Fig. 4F-right, Supplementary Fig. 4B**).

**Figure 4:**
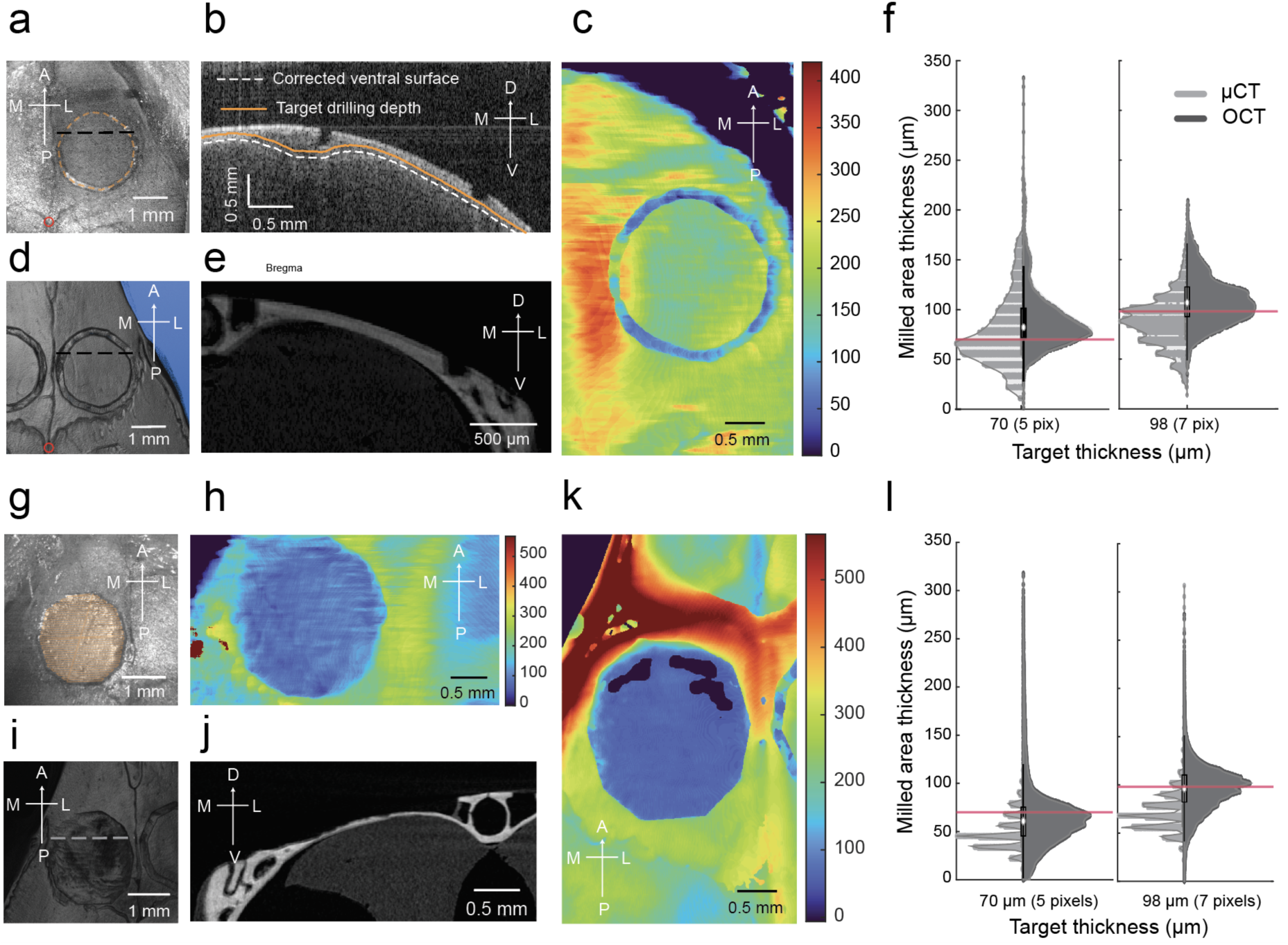
Robotic milling of mouse skull using estimated ventral surface as reference: **(a)** Reconstructed top view of OCT scan after milling circular trench on a mouse skull. **(b)** Cross-sectional view of the milled skull along the dashed line shown in a. **(c)** Pseudocolor plot of measured skull thickness after milling of circular trench in a. **(d)** Reconstructed top view of µCT scan of the same mouse skull shown in **a**. **(e)** Cross-sectional view of the milled skull as shown in **e**. **(f)** Violin plots of measured thickness of the skull along the milled trench path with target remaining thickness of 70 µm (left plot) and 98 µm (right plot), n = 4 mice. Red lines indicate targeted milling depth. **(g)** Reconstructed top view of OCT scan after milling over a circular area to thin the tissue down to 98 µm. **(h)** Pseudo color plot of the skull thickness measured via OCT scanning in the milled skull area shown in **g**. **(i)** Reconstructed top view of µCT scan of the same mouse shown in **g**. **(j)** Cross-sectional view of the milled skull as shown in **g**. **(k)** Pseudo color plot of the skull thickness measured via µCT scanning in the milled skull area shown in **g**. **(l)** Violin plots of the measured thickness of the skull in the milled circular area with target remaining thickness of 70 µm (left plot) and 98 µm (right plot), n = 4 mice. Red lines indicate targeted milling depth. A - Anterior, P - posterior, M - Medial, L - Lateral, D - Dorsal, V - Ventral. Red circles in **a** and **d** indicate Bregma.

The OCT measurements were cross-validated using µCT scanning of the same samples, with µCT providing a higher signal-to-noise ratio and slightly better resolution (11 µm both in lateral and axial directions). The µCT scan revealed rectangular profile trenches created by the flat end mill along the craniotomy (**Fig. 4D** and **E**), with the median thickness of bone remaining across 4 craniotomies performed being 68 µm with IQR: 55 µm-99 µm when milling was commanded to leave bone of thickness 70 µm. The remaining thickness had a median of 88 µm and IQR: 70 µm-102 µm when milling was commanded to leave the bone of thickness 98 µm. Showing that the ventral surface estimations were adequate to enable removal of the skull bone to the desired thickness.

We next performed robotic skull thinning procedures (8 procedures, n = 4 mice) where the skull within a 2 mm diameter circular area centered at either ∼1.3 mm posterior and ∼2 mm lateral to Bregma or ∼1.8 mm anterior and ∼1.4 mm lateral to Bregma. The CV-Craniobot was set to uniformly remove bone, leaving 70 µm for half of the procedures and 98 µm thick bone above the ventral skull surface for the rest of the procedures. Such skull thinning operations are typically performed in experiments that are particularly susceptible to potential neuroimmune responses to skull removal [28], [29]. Both post-milling OCT and µCT scans reveal robust tracking to the desired milling depth across the entire area targeted for thinning (**Fig. 4G-J**). The median thickness of the bone above the ventral surface was measured to be 61 µm (IQR: 45 µm-75 µm) when milling was commanded to leave the bone thickness of 70 µm and 96 µm (IQR: 82 µm-110 µm) when milling was commanded to leave 98 µm. Measurements of the same skulls made using the µCT revealed a median thickness of 45 µm (IQR: 34-57 µm) for the 70 µm target and a median of 68.1µm (IQR: 57-79 µm) for the target thickness of 98 µm (**Fig. 4K** and **L, Supplementary Fig. 5**). A slight decrease in the measured thickness in the µCT scans was observed compared to the measurements taken with the OCT immediately after the procedure. Possibly, this is due to the dehydration of the soft bone tissue or the trabecula.

These experiments confirmed that the information provided by non-destructive OCT imaging can be reliably used to accurately and precisely remove bone to a specific thickness in a robotic fashion.

### Computer vision-guided robotic milling enables rapid and precise craniotomies

We next sought to determine the maximum thickness of the bone from the ventral surface that can remain intact for a craniotomy to be successfully completed. Ideally, bone removal can be performed down to the ventral skull surface for complete bone removal. However, this operation might result in inadvertent contact with the underlying brain and dura due to potential errors in the model estimate. Experienced manual craniotomy surgeons typically aim to mill down to the soft part of the bone tissue, the trabecula, where the bone is loose enough to be fractured and removed. CV-Craniobot was programmed to perform 2 mm diameter circular craniotomies at the parietal bone (centered at ∼1.3 mm posterior and ∼2 mm lateral to Bregma) and frontal bone (centered at ∼1.8 mm anterior and ∼1.4 mm lateral to Bregma). The milling depth was adjusted to leave the bone thickness varying from 28 µm to 84 µm above the dorsal surface (**Fig. 5A**).

**Figure 5:**
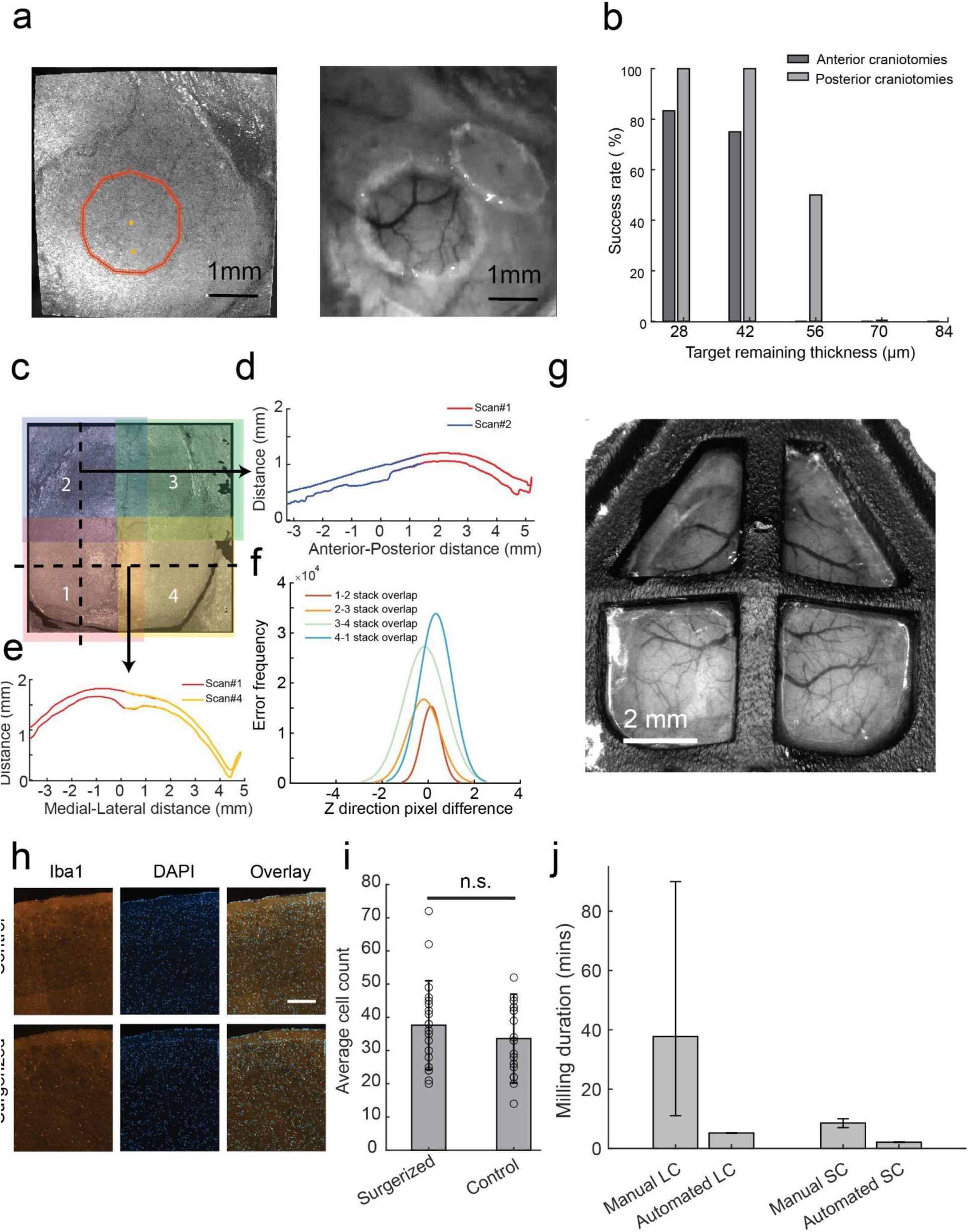
Automated craniotomies using CV-Craniobot: (a) *Left:* Top view of reconstructed OCT image of the surgery area, with path of circular craniotomy highlighted. *Right:* Photograph of the same location shown on the left taken after automated removal of the bone island. **(b)** The success rate of complete bone removal at different thicknesses of bone remaining on top of the ventral surface by the CV-Craniobot. **(c)** Top view of Computationally stitched reconstructed images from 4 OCT scans. The image from each scan is coded with a different color. **(d)** Sagittal cross-section of the stitched scans along the vertical dashed line shown in **c**. **(e)** Coronal cross-section of the stitched scans along the horizontal dashed line shown in **c**. **(f)** Alignment error during stitching between each two neighboring scans. **(g)** An example image of a mouse implanted with a multi-planar faceted cranial window covering the whole dorsal cortex was taken 21 days after robotic surgery using the CV-Craniobot. **(h)** Representative images of brain slices stained for Iba1 and DAPI of the brain directly underneath the craniotomy and control samples. The scale bar represents 200 microns. **(i)** Average microglia cell counts in a 700 x 900 µm² area under the milled skull area for the test samples and similar areas for the control samples. **(j)** Average time to complete craniotomy using the CV-Craniobot and comparison with human surgeons. Small craniotomies (SC): n = 5 procedures using the robot, n = 5 manual procedures performed by 2 experimenters. Large craniotomies (LC): n = 2 procedures using CV-Craniobot, n = 7 manual procedures performed by 3 experimenters.

It was observed that on the parietal bone, programming the CV-Craniobot to leave 28 and 42 µm (2 and 3 pixels in OCT image coordinates) of bone remaining intact over the dorsal skull surface resulted in a 100% success rate in complete bone island removal after a single round of milling (**Fig. 5B)** (n = 10 craniotomies attempted). The success rate of bone removal dropped to 50% (n = 10 craniotomies attempted) when leaving 56 µm of the bone thickness (n = 10 craniotomies attempted) and 0% when the CV-Craniobot was programmed for the remaining thickness of 70 µm and 84 µm (n = 10 craniotomies attempted for each target thickness).

Similarly, when performing craniotomies in the frontal bone, we found that programming the CV- Craniobot to leave 28 µm (2 pixels in PCT image coordinates) of bone remaining intact over the dorsal skull surface resulted in an 83% success rate in the complete bone island removal (**Fig. 5B,** n = 10 craniotomies attempted). The success rate of bone removal dropped to 75% (n = 10 craniotomies attempted) and 0% when (n = 10 craniotomies attempted) when the CV-Craniobot performed the craniotomy, leaving 42 µm and 56 µm of bone remaining above the dorsal skull surface. Similar to the parietal bone craniotomies, no craniotomies with a target remaining thickness of 70 µm and 84 µm resulted in a successful bone removal after a round of milling (n = 10 for each target depth). Compared to craniotomies performed over the parietal bone, the success rates at each milling depth were lower, possibly due to inaccurate estimation of the dorsal skull surface by the U-Net model or high variation in the refractive index of the bone. Note that the U-Net model training and the refractive index estimation were done mainly on parietal bone. Nevertheless, these results demonstrate the feasibility of single-shot rapid craniotomies, resulting in complete bone removal using the CV-Craniobot.

The size of the craniotomy that could be performed after scanning a single region of interest was limited by a 5 X 5mm lateral field of view (FOV) of the OCT imaging system. One of the advantages of automating craniotomies is the ability to perform very large craniotomies covering much of the dorsal skull surface [3], [5], [8], [9], [11], [18], [30], [31]. To perform such large craniotomies, the mouse skull surface was scanned across four overlapping FOVs, and the dorsal and ventral surfaces of the skull in each scan were estimated using the U-Net model.

Then, the surfaces were computationally stitched together (see **Methods**) to create composite dorsal and ventral surfaces of the whole dorsal skull (**Fig. 5C**). The stitching algorithm successfully aligned surfaces from the four scans. Neighboring surfaces were mostly aligned perfectly, generating a complete 3D representation of the dorsal and ventral surfaces of the mouse dorsal skull (**Fig. 5D** and **E**). Limited misalignment between the neighboring scans was observed, which is within the limits of the imaging system resolution (**Fig. 5f**). As a demonstration, we used the CV-Craniobot to perform a large craniotomy, removing over 52 mm^2^ of the skull surface over the dorsal cortex, encompassing an area over the primary and secondary motor, the somatosensory, association retrosplenial and visual cortices across both hemispheres to implant a multi-planar faceted cranial window [32] (**Fig. 5G**).

To assess the safety of the CV-Craniobot for automated craniotomies, we evaluated if the craniotomy resulted in an acute inflammatory response to the brain due to inadvertent contact of the milling tool with the soft brain tissue. The CV-Craniobot was used to perform 2 mm diameter craniotomies over the cerebral cortex with the target remaining thickness of 48 µm (centered at 2.6 mm medial to midline, 1.8 mm anterior to Bregma). All procedures resulted in successful bone detachment. The brain was covered with a gauze pad soaked in sterile saline (0.9% sodium chloride), and the animal was maintained under anesthesia for 60 minutes, following which animals were transcardially perfused and brain dissected for histological analysis. We stained the fixed brain slices for Ionized calcium-binding adaptor molecule 1 (Iba-1), a marker for activated microglial cells (**Fig. 5H**). A 700 X 900 µm area of the brain directly underneath the craniotomy had 38 ±*13* cells. The same area in the corresponding contralateral side had 34 ± 13 cells. There was no significant difference in the average number of IbA-1 positive cells among the two groups (**Fig. 5I**) (n = 10 samples, from under the surgerized areas and eight control samples (p=0.31, two-sample t-test). Thus, controlled bone removal using the CV- Craniobot results in no significant acute inflammatory response in the underlying brain tissue.

The performance of the CV-Craniobot was further assessed by comparing the speed of craniotomy procedures enabled by the CV-Craniobot to multiple manual surgeries performed in our laboratory. In contrast to manual craniotomy procedures, the CV-Craniobot needs to perform the OCT scan, followed by U-Net estimation of the dorsal and ventral skull surfaces.

Scanning each FOV took 44 seconds on average, while the U-Net surface estimation and milling path planning took approximately 2 minutes for each FOV. The CV-Craniobot is programmed to mill the bone at a maximum 6 mm/minute feed rate. Thus, the CV-Craniobot can consistently perform 2 mm diameter circular craniotomies in 2 minutes compared to the average of 8 minutes taken by human surgeons (**Fig. 5J**, n = 6 procedures completed by 2 surgeons), a 75% reduction in time. Surgeons included both experts (> 50 craniotomy procedures) and novices. The CV-Craniobot also performed large craniotomies across the dorsal cortex (**Fig. 5I**) in 5 minutes, an 85% increase in speed as compared to the average time taken during manual surgeries (34 minutes std: 23 minutes, n = 3 surgeons, 7 procedures). These results highlight the capabilities of the CV-Craniobot to perform precise and rapid craniotomy procedures in mice.

## DISCUSSION

We demonstrated that the CV-Craniobot could perform repeatable, precise, safe, and fast craniotomies on the dorsal mouse skull. Comparing milling durations, the CV-Craniobot achieved small craniotomies twice as fast as the previous version and was four times faster than other automated craniotomy systems in performing small circular craniotomies [15], [18], [33]. Minimizing the milling duration not only enhances the throughput of the procedure but also potentially increases the animal’s survival chance by reducing the duration of disturbance in the brain environment [34], [35]. Using OCT scanning to estimate skull thickness along the milling path eliminates the need for user intervention in specifying the milling depth, making the robot’s performance less prone to human errors.

Computer vision has previously been used to automate delicate neuroscience experiments such as image-guided patch clamping [36], [37], [38] and microinjection [39], [40], [41]. Integration of machine learning models for object recognition has further extended these efforts both by increasing the accuracy and specificity of object detection [42], [43] in a wide range of imaging conditions and providing variability in the types of objects and anatomical features that can be detected and manipulated robotically. Here, we harness deep learning methods to precisely and rapidly estimate skull morphology from OCT images to automate craniotomies in rodents. In the future, our approach for rapid 3D morphology detection using machine learning on non-destructive imaging for surgical operations could be harnessed in clinical settings for automating delicate microsurgical procedures.

The CV-Craniobot enables a unique preparation method of reducing skull thickness to a set value. Although skull thinning preparations have been performed and demonstrated in the past [18], [29], this is the first time that the skull can be thinned to a “uniform” thickness. This can improve imaging quality through the thinned window while minimizing disturbance in the brain environment. OCT scans provide valuable information beyond estimating the skull’s dorsal and ventral surfaces. The speckle variance in OCT images can map micro-vasculatures in the imaged tissue [44], [45]. This can potentially be used to study the brain, optimize the device implantation process, or enhance craniotomy path planning.

CV-Craniobot was designed with modular bays, each dedicated to performing specific tasks. This design allows for the easy integration of new functionalities by adding additional bays. For example, an electrode insertion bay could be incorporated to enable precise microelectrode insertion, with control over the insertion angle based on OCT scans of brain tissue. Similarly, bays for assisted implantation procedures—ranging from implant placement to using additive manufacturing methods to 3D-print neural interfaces on the animal [46]—can also be envisioned.

The 3D skull reconstruction performed by the CV-Craniobot was not as accurate as μCT scans due to the U-net underestimation of skull thickness in some thicker areas like the central sinus and frontal bone. Other factors include variability in the refractive index of skull tissue and limitations in the model used to correct optical distortions, particularly in extreme lateral regions of the scan [21], [22]. The accuracy of surface estimation can be enhanced by improving the trained model and deploying more sophisticated scan distortion correction techniques. Despite these inaccuracies, the experiments demonstrated that the current estimations were accurate enough to provide a functional assessment of each surface’s depth.

## METHODS

### CV-Craniobot hardware construction and design

The CV-Craniobot hardware comprises of four linear actuators: two compact, motorized translation stages with a travel range of 50 mm (MTS-50 Z8, Thorlabs Inc.) for moving the dental driller and the OCT scanner along the Z-axis, a 300 mm range stepper motor linear translation stage (LTS300C, Thorlabs Inc.) for moving the custom-made stereotax between the OCT stage and the milling stage and adjusting the X-position of the sample, and a DC-servo motor linear translation stage with a range of 30 mm (M30X, Thorlabs Inc.) for moving the stereotax in the Y direction. The custom-made aluminum stereotax, designed similarly to the Craniobot [30], is mounted on the X and Y linear stages, which are assembled orthogonally using a custom-designed connector made from 3/8" thick machined 6061 aluminum sheets. The Z-actuators are mounted on 20" construction rails (50 mm x 75 mm, XE5075L20, Thorlabs Inc.) using custom-made aluminum connectors. The OCT scanner and the USB digital microscope are mounted on the Z-actuator using a custom-designed, 3D-printed holder. The milling procedure employs an electric dental drill (RAMPOWERDIGITAL45, RAM Products, Inc.) mounted on the Z-stage with another custom-designed, 3D-printed holder.

<u>The OCT scanner:</u> OCT scanning of the mouse skull during surgeries was performed using a commercial spectral domain OCT scanner with a center wavelength of 1310 nm and a bandwidth of 60 nm (OQ Stratascope 1.0, LUMEDICA INC) and equipped with a custom objective having a fixed working distance of 1”. The native software package (Lumedica OQ LabScope, LUMEDICA INC) was used to manage the scanning process, perform primary calculations on the raw signals, and generate cross-sectional B-scans.

<u>Software communication and control:</u> The CV-Craniobot control software was developed in LabVIEW (National Instruments Inc.). The motorized stages were connected to the control computer via USB serial communication, linking the computer to each linear stage’s controller. The linear stage controllers used a built-in PID (proportional, integral, derivative) algorithm to execute position commands and communicated with LabVIEW through .NET controls provided by their native software (Kinesis®, Thorlabs Inc.) (see **Supplementary Fig. 6** for an illustration of the communication diagram). The CV-Craniobot controller software communicated with the OCT scanner’s native software through a virtual serial port (virtual serial port driver, version 11, Electronic Team Inc., 2023). Multiple scripts for calculating the milling path based on the OCT scans of the sample were written in MATLAB (MATLAB ® - MathWorks), which were called and executed through the LabVIEW interface for MATLAB.

### CV-Craniobot calibration

To perform automated milling based on the OCT scans of the sample, it is necessary to determine the relationship between the coordinates of each 3D point in the OCT scans and the position of each motorized stage. After the OCT scan distortion removal process, the transformation operator was found using Normalized Direct Linear Transformation (NDLT) [47]. To correlate the 3D position of the driller’s tip (X-stage, Y-stage, and driller’s Z-stage positions) with the 3D space of the OCT image, nine holes were automatically drilled in modeling clay using the dental driller, with its position controlled and recorded by the linear stages. The drilled sample was then transferred to the OCT stage and imaged by the OCT scanner. After normalizing the points’ coordinates, the relationship between the recorded drilling position for each hole and the position of the center of the drilled hole on the OCT scan was extracted by finding the affine transformation matrix (calibration matrix) relating these two spaces. To ensure a robust solution and increase accuracy, the pilot points were designed so that no three points were collinear and no four were coplanar [30]. The resulting calibration matrix and the position of the robot stages during OCT imaging were then saved for further use.

Processing of the OCT scans and all related calculations to find the calibration matrix was performed using custom MATLAB scripts with the MATLAB Image Processing Toolbox. All MATLAB scripts, along with the custom-built GUI, were embedded in a LabVIEW VI designed specifically for the calibration procedure. The script also contained a module to annotate the edges of the OCT’s “top-view” field of view in the live images from the USB microscope (MS100 portable USB microscope, Teslong Technology) mounted along with the OCT scanner. The annotated area in the live video from the USB camera was used to ease the sample adjustment process under the OCT scanner prior to automated craniotomy.

### Non-telecentric distortion correction

OCT scans were corrected for non-telecentric scanning distortions prior to being used for the dorsal and ventral surface estimation by the U-net model.

Fan distortion, i.e., non-telecentric scanning distortions, were modeled by radial and tangential distortions in Z cross-sections of the OCT scan. To find the distortion parameters in the resultant XY image, the imprint of a piece of woven wire 120 mesh on modeling clay was imaged using the OCT scanner at different distances from the scanner with increments of 75 μm. For each batch of B-scans (one volume scan), the XY image was generated as the Z cross-section of the scan at Z equal to the mean clay surface Z-value. The resultant XY images for each B-scan batch are shown in **Figure 2B**. The edges of the mesh’s imprints were detected using the MATLAB image processing toolbox function (edge) using the “Canny” method, labeled, and stored as separate image components using MATLAB’s function (bwlabel). Each detected edge represented a straight line that was deformed due to the image distortions.

<u>Radial distortion correction:</u> The division model [48] of radial distortion was used to model and correct for radial distortion in the XY image. The division model can express high distortion in low order, sufficient for commonly seen distortions in cameras [49], [50], [51]. Hence, the model’s first order was used to express the image’s distortion. Specifically, Wang et al. [51] method was used, where each line was fitted on a circle, and the radial distortion parameters were found as the solution to a system of linear equations relating the parametric equation representing a straight line after distortion and the equation representing the distorted lines fitted on a circle. The circle fit to each deformed line was found using a custom MATLAB script developed previously [52], which uses the direct least-squares fitting method to find the best-fit circle. The system of linear equations for all lines was solved using a custom MATLAB script, and the radial distortion parameters (distortion center and first-order radial distortion coefficient) were extracted for the XY image. The same procedure was repeated for all OCT volumes, and for each Z, the radial distortion parameters were calculated. The resultant radial distortion parameters were calculated as the mean of these values across different Z cross-sections. The extracted parameters were used to recalculate the position of each point in the 3D scan.

<u>Tangential distortion correction:</u> The Brown-Conrady model, which represents both tangential and radial distortion, was employed to correct for tangential distortion in the XY image for each Z value [53]. After correcting the image for radial distortion, the first-order tangential portion of the Brown-Conrady model was used to model the distortion in the image. Tangential distortion parameters were calculated by first reconstructing the undistorted lines as lines parallel to either of the two perpendicular lines crossing the center of radial distortion in the same position as the current distorted lines. Then, the parametric equations for each line were reconstructed using Brown’s model. The parametric equations were compared to the distorted lines detected in the image, and the distortion parameters were extracted. The average of the tangential distortion parameters over different Z values was chosen as the final tangential distortion parameter. After reconstructing the corrected scan, the empty pixels were filled by interpolation using a custom function (ndnanfilter) written in MATLAB [54].

To assess the success of the image correction algorithm in reducing telecentric distortion in the XY-plane of the OCT scans, the root mean square error (RMSE) of each point on an X or Y line in the distorted image was calculated. The error for each point on a line was defined as the distance between that point and the corresponding point on the reference line (undistorted form of the line). The error was calculated for all XY images from Z = 72 to 337 pixels (the usable range for imaging through the mouse skull). The same procedure was applied to the corrected image, and the RMSE was calculated for each detected line. The entire process was performed using a custom MATLAB script.

### OCT top-view image generation

A custom MATLAB function was used to reconstruct the top-view image of the scanner’s FOV, enabling the user to choose the position of the milling path on the animal skull. The function reads the OCT scan from the hard drive and corrects the telecentric distortions in the scan. The top-view image was reconstructed by summing the intensities for all Z values for each X and Y, replicating what a 2D camera would do to generate a top-view image of the sample. The image was further refined using MATLAB’s built-in image processing toolbox function ("localcontrast") to adjust the image’s contrast, setting the maximum and minimum intensity range of the image within 3 standard deviations from the mean intensity value.

### U-Net model training

A U-Net model was used to perform semantic segmentation of each B-scan, which was then post-processed to extract the dorsal and ventral boundaries. The model was trained for 100 epochs using the Adam optimizer [55] with a learning rate of 4e-4. For our loss function, we used the region-based Tversky loss [56], [57], finding that it produced more structurally coherent predictions—especially around borders—compared to distribution-based criteria such as binary cross entropy. We implemented training augmentations to improve the model’s generalization to novel samples which included a random horizontal and vertical flip (p=0.5), as well as a random vertical translation (p=0.5). Model performance was quantified using the pixel-wise accuracy and Intersection-over-Union (IOU) metrics on out-of-fold samples within a K-Fold cross-validation scheme (k=5). The model was trained on 300 annotated live mouse skull B-scans. Four scans from the same FOV were averaged into a single high-quality scan for each FOV to enhance the quality of the manual annotations. These high-quality scans were then selectively annotated in the corresponding B-scans. For the training, the annotated areas were overlaid onto the raw image from one scan. A custom MATLAB script was used to prepare the training set.

### Dorsal and ventral surface reconstruction

A compiled custom MATLAB script was used to estimate the positions of the dorsal and ventral surfaces. As explained above, the script first reads the OCT stack and corrects it for distortions. Then, it smooths the scan in the Y direction using a 3D median filter (a built-in MATLAB function) with a window size of 15 pixels in the Y direction. Next, the script uses the trained U-net model to infer the pixels containing the mouse skull in each B-scan. The dorsal surface is calculated as the largest gradient in the Z direction in the resulting binary cross-section image from the U-Net model (from 0/no skull to 1/skull), and the ventral surface is found as the lowest gradient in the Z direction (from 1/skull to 0/no skull). The calculated dorsal and ventral boundaries of the skull in each B-scan are then accumulated to generate 3D surfaces of the dorsal and ventral areas of the skull. The surfaces are further smoothed using a two-dimensional median filter with a window of 5 pixels in the X direction and 10 pixels in the Y direction. Finally, the script adjusts the orientation of the surfaces for ease of calculation in the subsequent steps and saves the generated surfaces along with the top-view image of the OCT field of view as a .mat file on the hard drive. The script is compiled into an executable file using MATLAB Compiler to facilitate the use of the trained model by LabVIEW VI and reduce the complexity of system configuration for future system replication.

### Optical distortion (refractive index) correction

To correct the OCT images for optical distortion, every surface imaged through a material other than air should be corrected for depth. This is accomplished by dividing the thickness of the medium by its refractive index[21]. Hence, the corrected depth of the ventral surface was calculated as follows:

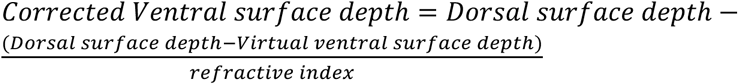

After estimating the dorsal and ventral surfaces of the mouse skull using the U-net model, the depth of the ventral surface at each point was corrected using the above equation with a custom MATLAB script.

### Milling path computation

Non-axial (X and Y) positions of the milling pilot points with respect to the center of the craniotomy were provided to the system in millimeters as user input in a text file format following the control software prompt. The milling depth for each pilot point was automatically calculated through a custom MATLAB script embedded in the CV-Craniobot’s control VI in LabVIEW as follows:

After completing the OCT scanning, a top view of the sample was generated from the OCT scans. The user was prompted to annotate the center of the milling axis and a point on the negative Y axis on the top-view image. The input pilot points were then rotated, centered based on the input path center and orientation, and scaled to match the OCT magnification (each millimeter is approximately 111 pixels in the OCT transverse direction). The resulting milling path was overlaid on the OCT top-view image for the user’s confirmation. To ensure the milling pilot points were sufficiently close to capture the sample height changes along the milling path, a custom MATLAB function was used to linearly interpolate between the original pilot points to keep their distance below 100 μm. The interpolated points were then added to the original set of milling pilot points. The milling depth was calculated using different methods depending on the input milling mode (constant remaining thickness milling or constant trench depth milling). For each pilot point, the depth was determined by first referencing the depth of the reference surface at that point. For constant trench depth procedures, the reference surface was the dorsal surface, while for the “constant remaining thickness” mode, it was the skull’s ventral surface. The user was then prompted to input the desired milling depth or remaining thickness (based on the milling mode). The depth of the milling path was calculated as an offset from the reference surface (positive offset for “constant remaining thickness” and negative for “constant trench depth”).

In the final step, the 3D coordinates of the milling pilot points were transformed from OCT 3D space to the linear stage’s coordinate space using the calibration matrix and the relative position of the sample during OCT scanning with respect to the calibration sample during the calibration process:

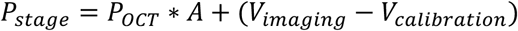

Where 𝑃_*stage*_ is the milling pilot points in the stage’s axis, 𝑃_*OCT*_ are the pilot points coordinates in the OCT space, A is the calibration matrix, 𝑉_*imaging*_ is the position vector of the stereotax and the (X and Y) and the OCT scanner (Z) during the sample’s scanning process and 𝑉_*calibration*_is the position vector of the stereotax while imaging during the calibration process. The generated milling pilot points were saved as a text file on the hard drive and passed to the software controller to perform milling.

### Surgical procedures

All animal experiments in this study were conducted under approved protocols by the University of Minnesota Institutional Animal Care and Use Committee (IACUC). Mice from three strains (Ai162/Cux2-cre, Ai162, and Thy1-Gcamp6f) of both sexes and ages ranging from 10 to 56 weeks were used. Prior to surgery, the mice were anesthetized with 1–5% isoflurane (Piramal Critical Care Inc., Bethlehem, PA) and were administered Buprenorphine (Par Pharmaceutical, Chestnut Ridge, NY) at 1 mg/kg and Meloxicam (Dechra Veterinary Products, Overland Park, KS) at 1–2 mg/kg. Buprenorphine was replaced by slow-releasing buprenorphine (Buprenorphine SR LAB, Zoopharm, Windsor, CO) at 1 mg/kg, and Meloxicam was replaced by Dexamethasone at 2 mg/kg for survival surgeries. The mouse scalp was shaved and cleaned, and the animal was transferred to the CV-Craniobot’s stereotax, which maintained anesthesia with a continuous supply of isoflurane in oxygen. The scalp area was sterilized using standard aseptic techniques, and the scalp and fascia were manually removed with autoclaved surgical instruments. The mouse skull was dried with autoclaved cotton swabs. At the initiation of the robotic craniotomy procedure, the mouse was transferred to the OCT imaging bay for scanning.

After scanning and path planning, the mouse was automatically transferred to the milling bay, where a craniotomy was performed. During milling, the skull was constantly irrigated with sterile saline to remove bone dust and prevent tissue damage from the heat generated during milling. An additional OCT scan was performed for assessment purposes at the end of the craniotomy procedure. The animal was euthanized after removal from the stereotax after acute surgical procedures. During the survival surgery, a sterilized cranial implant was fixed on the skull with dental cement (S380, C&B Metabond, Parkell Inc.) after bone removal. Once the surgery was done, the mouse was transferred to a clean, heated cage for monitoring until full recovery. The animal’s state was monitored for up to three days post-procedure.

### Automated craniotomy using the CV-Craniobot

Mice were head-fixed on the custom stereotax, and the scalp was removed. Then, CV-Craniobot control software guided the user through multiple steps via user prompts for inputs. The user could choose the milling mode - i.e., using the dorsal skull surface as a reference or ventral skull surface as a reference, and specify the craniotomy’s size, shape, and location.

User-initiated inputs guided the affixed mouse first to the OCT scanner bay, where the head of the mouse was adjusted via visualization using the targeting camera (MS100 portable USB microscope-Teslong Technology) to ensure that the area of the desired craniotomy was within the OCT imaging FOV. The height of the OCT scanner with respect to the skull was adjusted using the live B-scans in OCT’s native control software. The CV-Craniobot next scanned the skull using the OCT scanner and generated a top view of the reconstructed OCT scan image.

Next, the user specified the orientation and location of the milling path by annotating the top view image, after which the CV-Craniobot computed the required milling path. In the next step, the CV-Craniobot moved the stereotax to the milling bay to perform a robotic craniotomy. The milling procedure was carried out using an electric dental drill operating at a speed of 20000 to 30000 rpm, equipped with a miniature square-end mill with a diameter of 200 μm (Harvey Tool Inc., 13908). A complete automated circular craniotomy procedure is illustrated in **Supplementary Video 2**.

### Constant trench depth milling on a live mouse skull

A 20-week-old male Thy1-Gcamp6f mouse was used to perform multiple constant-depth craniotomies. Using the CV-Craniobot, five horizontal lines perpendicular to the central suture orientation were milled on the left hemisphere of the mouse skull. Each line had a different trench depth, ranging from 42 to 98 μm. Similarly, five vertical lines with varying trench depths were milled on the right hemisphere of the mouse skull. After each craniotomy, the mouse was automatically transferred back to the OCT stage for scanning. These scans were later used to assess the milling depth in each procedure.

### Constant remaining thickness milling and Skull thinning procedure

Five mice (four females and one male), aged 10 to 15 weeks and from various strains (Ai162 and Ai163/Cux2- cre), underwent two circular craniotomies and two circular skull thinning preparations with different target remaining thicknesses. The position of each craniotomy varied between mice to assess the CV-Craniobot’s performance in different areas of the skull. The milled area was scanned for performance assessment at the end of each craniotomy procedure. After the experiment, the animals were euthanized, and their skulls were extracted and preserved in 4% PFA.

The skull thinning procedure involved constant remaining thickness milling along a specific milling path designed to cover the entire area of interest. The milling path was generated using a custom MATLAB script that calculated a raster milling path for any drown convex shape. The distance between each milling pass could be adjusted for different end mills. The distance between each pass was set to 80 microns for the experiments reported in this paper.

### Milling procedure assessment using the OCT scanner

To assess the performance of the automated system, the sample was returned to the initial scanning position (𝑉_*imaging*_) after the milling procedure, and an OCT scan was performed. The dorsal surface of the sample after milling was calculated by finding the maximum gradient in the Z direction for each X and Y. U-Net was not used to find the surfaces after milling due to its inferior performance compared to the gradient method in finding the depth of the milled trenches. A global offset between the dorsal surfaces before and after milling in non-milled areas was detected in some cases. This offset was partly due to the minor differences between the U-net and gradient thresholding methods in identifying the dorsal surface pixels. This height difference (dorsal offset) was calculated as the mode of the differences between the two dorsal surfaces.

The difference between the two dorsal surfaces (before and after milling) was measured along the milling path for constant-depth drilling. It was reported as the final milling depth after being corrected for dorsal offset. For constant thickness drilling, the distance between the milled dorsal surface and the ventral surface (corrected for dorsal offset) before milling along the milling path was used as the remaining thickness measure. The whole process was performed using a custom MATLAB script.

To assess the linear relationship between the targeted milling depth and the resultant milled depth, the measured thicknesses along the path against the target depth were fitted for all thicknesses using robust (“bisquare”) linear regression using a built-in function (“fitlm”) of statistics and machine learning toolbox in MATLAB, the coefficient of determination was also extracted as a property of the output model.

### Micro-CT scanning and processing

To cross-validate the OCT measurement, the samples were scanned shortly after the OCT scan with a micro-computed tomography scanner (H225 X- teck XT, Nikon PA, USA). The scans were taken with a resolution of 11 microns on all axes, and the FOV of the scans contained the entire mouse skull. To prepare the samples for μCT, each mouse was euthanized, and the skull was extracted and mounted on a Teflon pedestal compatible with the micro CT’s scanning stage. The sample was fixed on a mount compatible with the micro CT scanner using epoxy glue (DP100 Plus Epoxy Adhesive-3M) and was scanned with the same setting as our previous studies [27]. The X-ray images of the sample were then saved as a volume graphics file (.vgi) and subsequently saved as .tiff files using VGSTUDIO MAX 3.0 (Volume Graphics GmbH, Heidelberg Germany). The scans were then analyzed using a custom MATLAB script. Scans were smoothed using a 3D median filter with a window size of 3 pixels in all axes, followed by segmentation using MATLAB image processing toolbox function (“imbinarize”). The segmented binary image was then used to find the dorsal and ventral surfaces by finding the first and last pixel equal to one in the Z direction for all X and Ys.

### OCT-μCT scan comparison

The acquired OCT and μCT scans have different resolutions and fields of view. To compare the measured skull thickness through these different scanning systems, the dorsal and ventral surfaces of the sample were extracted using the methods explained above for each scanning modality. Then, the μCT scan was transformed using an affine transformation matrix to have the same orientation and magnification as the OCT scan. After the transformation, the thickness of the mouse skull was calculated for each modality by subtracting the depth of the dorsal surface from the ventral surface.

The affine transformation matrix for transforming the μCT scan was calculated using a custom script that finds the solution for the Matrix M as a solution to the linear equation P=Q*M.

P is the position of visible skull landmarks in the OCT scan, and Q is the position of the same landmarks in the μCT scan. The user manually annotated the landmarks in both scans.

### Bone removal success rate experiment

To assess the success rate of bone removal at different remaining thicknesses of the mouse skull under the milling path, multiple circular craniotomies with a diameter of 2 mm and varying target remaining thicknesses (28 to 84 μm) were performed on eight mice, aged 17 to 21 weeks. The group included two males and six females from various strains (Ai162/Cux2-cre, Ai162, and Thy1-GCamp6f). Each mouse underwent 3 to 8 craniotomies, totaling 10 for each target thickness. After automated milling, the experimenter tested the bone disk to verify if it was removable and recorded the result.

### Histology

The histology study was performed on a subset of mice (n = 5 mice, aged 7 to 23 weeks, comprising two males and three females from various strains (Ai162/Cux2-cre, Ai162, and Thy1-GCamp6f). On each mouse, a circular craniotomy with a diameter of 2 mm and a predefined remaining thickness of 28 μm was milled on the left hemisphere of each mouse skull using the CV-Craniobot. After milling, it was confirmed that the bone island could be removed. The animals were kept anesthetized on the CV-Craniobot’s stereotax for 1 hour after the milling procedure to allow for a potential neuroimmune response due to tissue damage during the milling process. Mice were recovered from the robot and transcardially perfused using methodologies we have previously used [58]. The dissected brains were sliced and stained using the same procedure as established previously [18]. The number of microglial cells in an area of 700 × 900 µm under the milling path was counted using a custom MATLAB script. Four tissue samples were used from each mouse to count immune cells under the milling path.

Microglial cells were identified by the co-occurrence of Iba-1 and DAPI staining in the same area. The contralateral hemisphere of the same mouse brain was used as the control sample to count the baseline number of microglial cells in the tissue.

### Image stitching

To perform larger craniotomies using the CV-Craniobot, a larger area needs to be scanned by the OCT scanner. To extend the OCT scanner’s FOV, after the user positions the sample to fit in the target area (target position), the CV-Craniobot moves the sample to four different predefined positions with respect to the target position and performs OCT scanning.

These scans collectively cover the entire area of interest. The CV-Craniobot software analyzes each scan separately to infer the dorsal and ventral surfaces. The software stitches the resulting surfaces together using the known relationship between the position of X, Y, and OCT Z stages and the center of the image in the OCT scan (a calibration matrix), and the position at which each scan was performed with respect to the target position to predict the position of each surface in the larger OCT space. The Z value of the surfaces in overlapping regions is estimated as the average height of the overlapping surfaces.

## SOFTWARE AVAILABILITY STATEMENT

The CV-Craniobot control software is available on request.

## DATA AVAILABILITY STATEMENT

Data included in this manuscript will be made available upon reasonable request.

## ACKNOWLEDGEMENTS

We thank Bonita Van Heel and Dr. Alex Fok for help with µCT imaging experiments. The µCT scanning was performed at the Minnesota Dental Research Center for Biomaterials and Biomechanics. We thank Colleen Forster at the Center for Translational Science Institute (CTSI) for her help with histology experiments. The CTSI is supported by the National Center for Advancing Translational Sciences of the National Institutes of Health Award Number UM1TR004405 and UL1TR002494. We also thank Dr. James Hope for fruitful discussions on using OCT imaging for this project.

## AUTHOR CONTRIBUTIONS

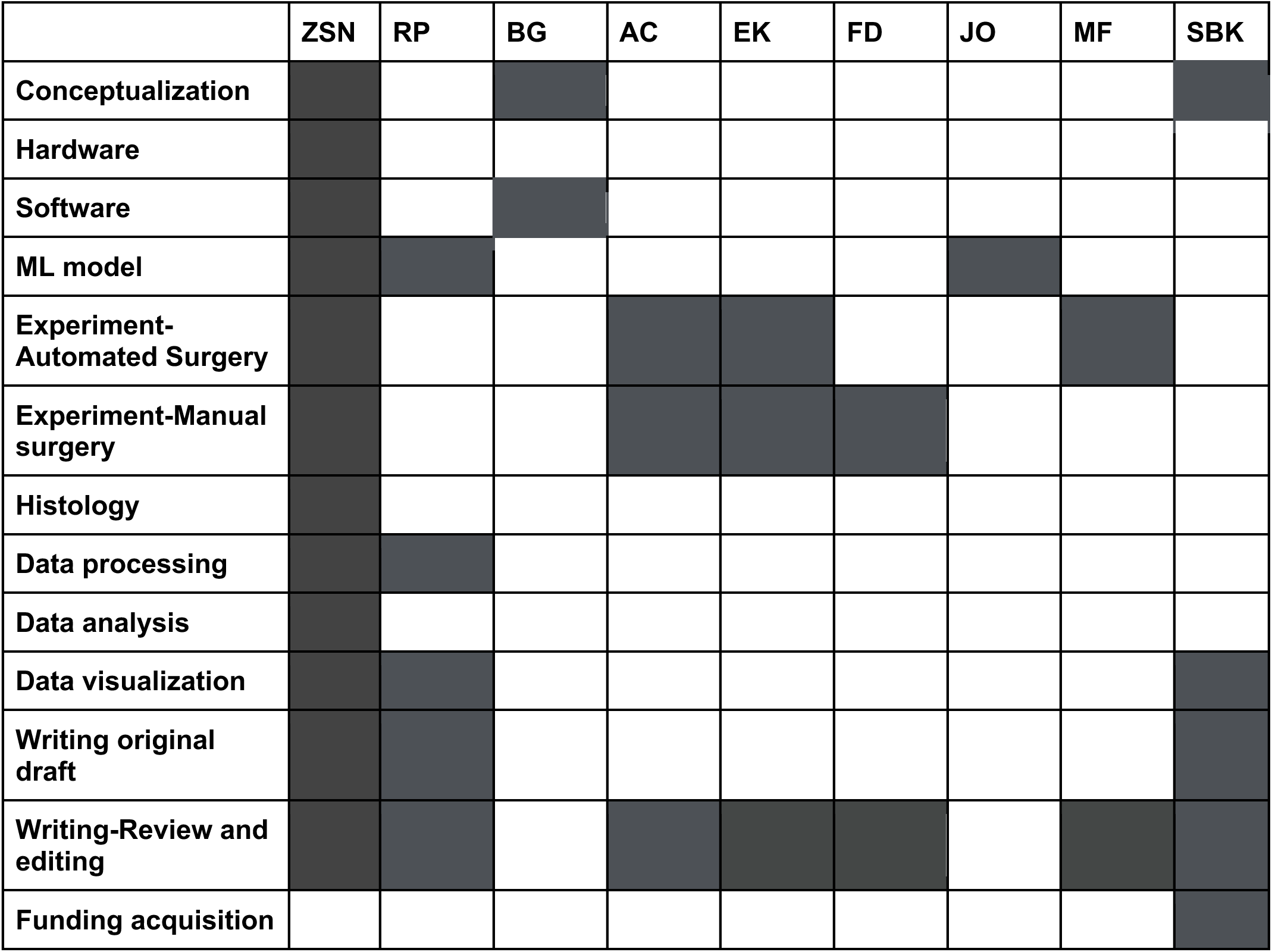

