## Supplementary Figures and Notes for "Computer vision guided rapid and precise automated cranial microsurgeries in rodents"

Navabi et al.

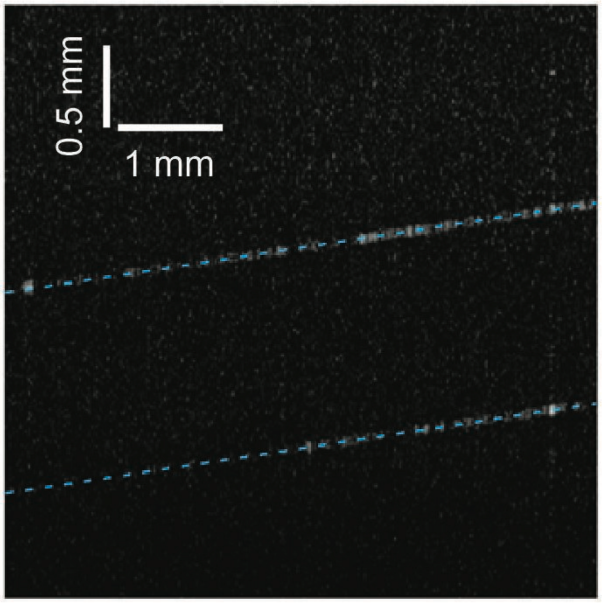


**Supplementary Figure 1:** OCT image of a glass cover slip. Blue dashed lines indicate the air-glass interface.

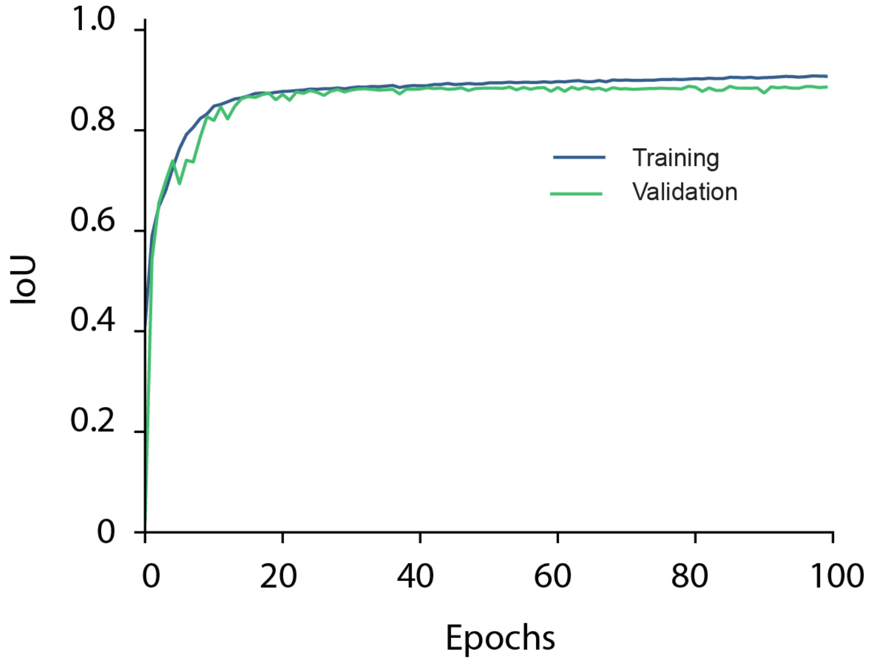

**Supplementary Figure 2:** The Intersection over the Union plot for the trained U-net model.

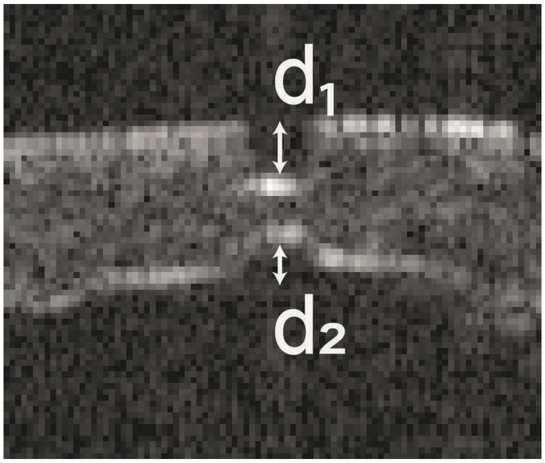


**Supplementary Figure 3:** B-scan image from OCT imaging taken after a partial burr hole is drilled into the skull. d1 - depth of drilling,  d2- depth of virtual hole with respect to the ventral surface.

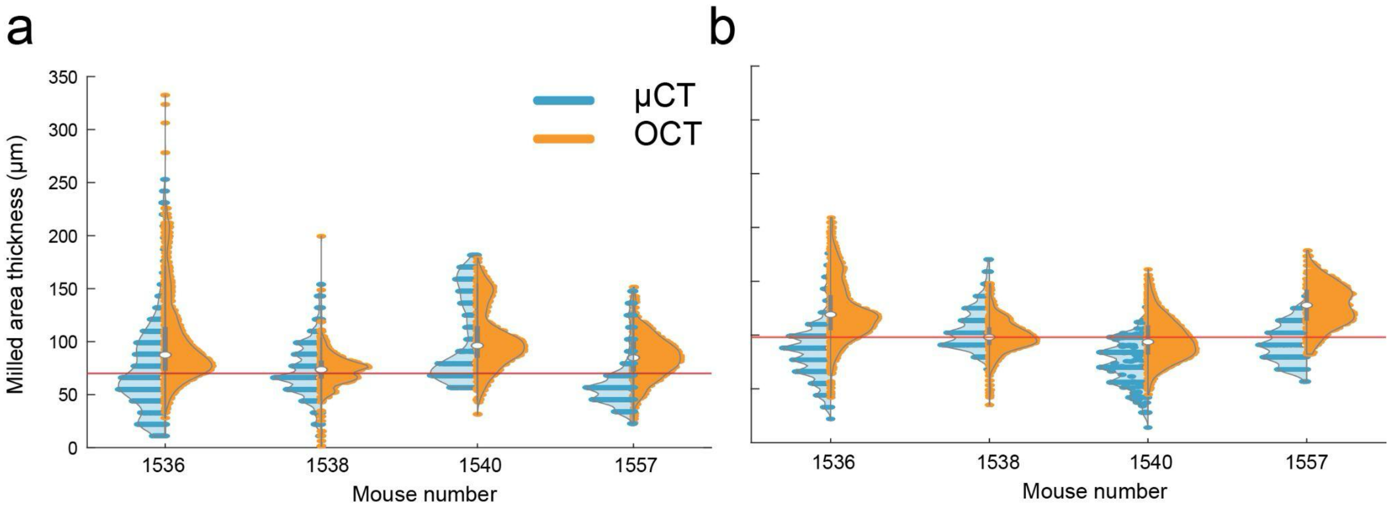


**Supplementary Figure 4:** Violin plots of the measured thickness of the skull in the milled trench path with the target remaining thickness of **(a)** 70 µm and **(b)** 98 µm shown for each mouse. Data is summarized in **Figure—4f** in the main manuscript. The red line indicates the target thickness.

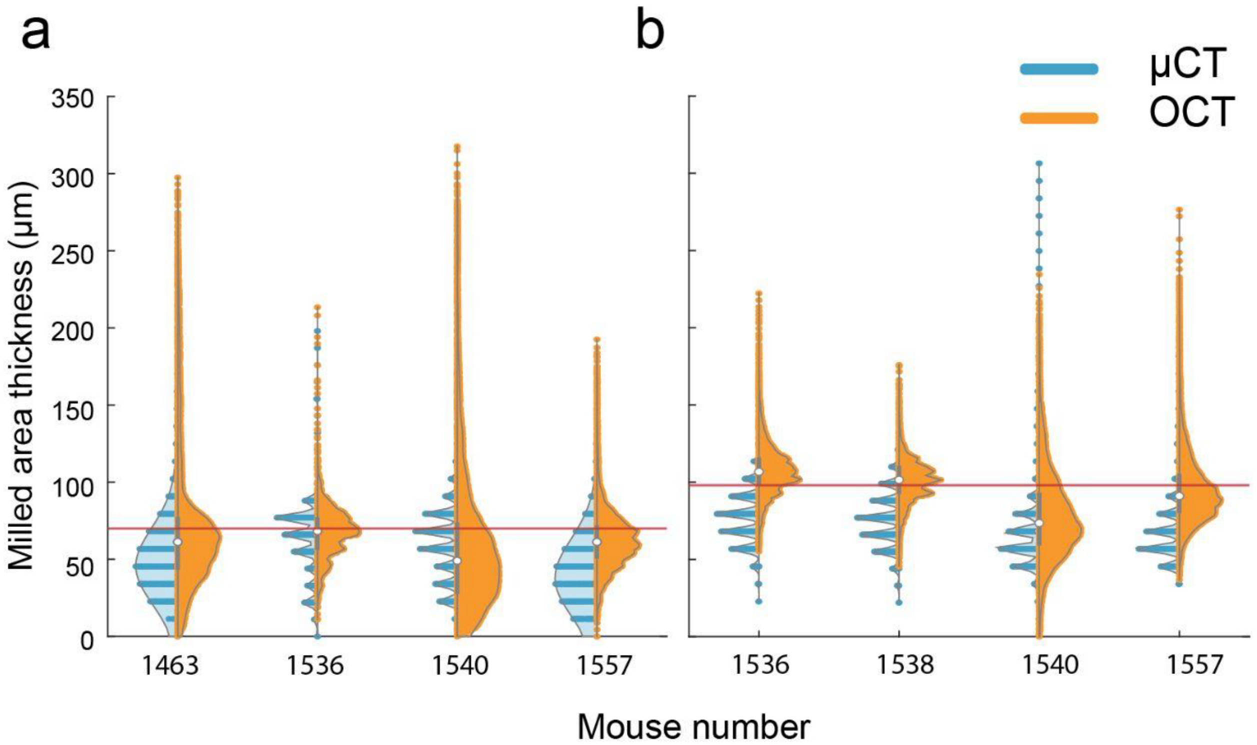


**Supplementary Figure 5**: Violin plots of the measured thickness of the skull in the milled circular are with target remaining thickness of **(a)**, 70 μm and **(b),** 98 μm (right plot) shown for each mouse. Data is summarized in **Figure—4l** in the main manuscript. The red line indicates the target thickness.

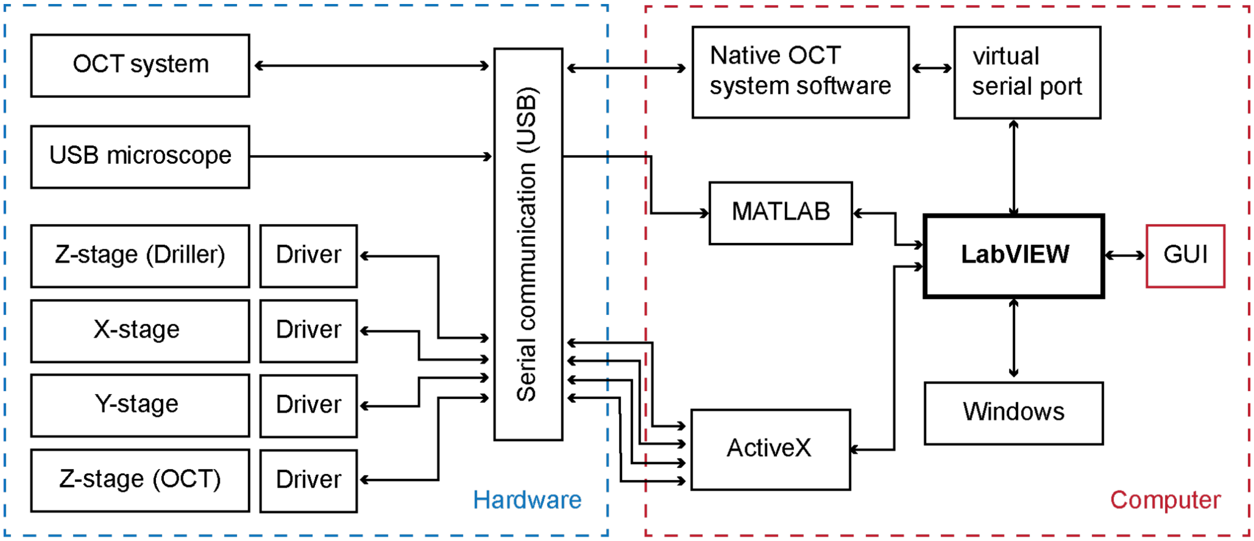


**Supplementary Figure 6:** Schematic diagram of digital communications between the CV-Craniobot hardware components.

**Supplementary Note 1: Measuring refractive index of mouse skull tissue using OCT imaging:** An OCT system determines scattering depth by measuring the difference in travel time between light returning from the sample and the reference arm. Due to the slower speed of light in bone compared to air, the light takes longer to return from the ventral surface of the skull, causing the dorsal surface to appear deeper in OCT images than it actually is [1], [2]. For instance, drilling a partial burr hole on the skull results in not only a lowering of the air-skull interface in the dorsal skull surface but also results in a raised virtual ventral surface in the drilled region (**Supplementary Fig. 3**). We used this property to systematically determine a nominal refractive index for the skull that could be used to correct for optical distortions [2], [3]. Partial burr holes with variable depths were drilled into the skulls of n = 8 mice (three females and five males, aged between 9 to 68 weeks from Ai162/Cux2-cre and Thy1-GCamp6f strains) during acute surgeries. The depth of the holes varied. Drilling was performed to ensure the largest depth was above the ventral surface of the skull. To clearly see the ventral edge of the mouse skull during these scans, two through-holes were drilled in each mouse. Air was gently blown through holes to separate the skull and brain and create a layer of air underneath the ventral surface to increase OCT image contrast. The depth of the drilled burr hole with respect to the air-dorsal skull surface interface, denoted as d1, and the depth of the virtual hole that appears in the drilled area of the skull, relative to the ventral skull surface in the OCT B-Scan, denoted as d2 are related via refractive index (RI) of the skull by the equation:

$$d_2 = d_1(RI - 1)$$

The RI was calculated by measuring $d_1$ and $d_2$ at multiple points inside the burr holes. As the refractive index varies due to changes in bone density and blood flow in the bone tissue, these measurements were made at multiple burr holes drilled at several locations on the dorsal skull. Burr holes where either d1 or d2 were less than 2 pixels (28 μm) were disregarded. The resultant histogram of the RI is shown in **Figure 2G**.
